# NanoporeDB: A Structural Resource Of Multimeric Protein Nanopores In Single-Molecule Sensing

**DOI:** 10.1101/2025.11.25.690617

**Authors:** Yuqian Liu, Zidong Su, Wenzhen Yang, Denghui Li, Jiawen Zhang, Yuning Zhang, Tao Zeng, Yong Zhang, Yuxiang Li, Guangyi Fan, Kailong Ma, Shanshan Liu, Xun Xu, Yuliang Dong, Zongan Wang

## Abstract

Protein nanopores are essential molecular gateways in biology and have inspired transformative technologies in biosensing and single-molecule sequencing. While this technology has transformed genomics and biosensing, the discovery of novel nanopore scaffolds remains limited due to the scarcity of experimentally resolved pore structures. Here, we present NanoporeDB, an open-access structural resource comprising over 6,600 high-confidence multimeric models across four representative pore types. Candidate proteins were systematically mined from large-scale datasets, including the AlphaFold Protein Structure Database, UniRef90, and MGnify90, and assembled using AlphaFold-Multimer and AlphaFold3. We performed membrane embedding, pore axis annotation, and constriction profiling to enable functional interpretation. NanoporeDB features an interactive web interface with 3D visualization and quantitative metrics such as insertion depth, tilt angle, and pore geometry. This resource provides a structural foundation for advancing nanopore-based molecular sensing, precision diagnostics, and synthetic biology.

## Introduction

Protein nanopores are versatile transmembrane channels that mediate selective molecular transport, sensing, and signaling in living organisms [1]. Their biomimetic application, leveraging tunable pore geometries, has inspired transformative technologies in biosensing, sequencing, and macromolecular analysis [2]. Among these, nanopore sequencing has emerged as a powerful single-molecule technique that enables direct, label-free analysis of DNA, RNA and proteins [3]. It relies on nanopores embedded in electrically insulating membranes (e.g., lipid bilayers) to form transmembrane channels with constriction zones, where voltage-driven translocation of analytes generates characteristic ionic current signals for real-time molecular identification and sequencing [4] (Figure 1A). Owing to its unique advantages in long-read capability, real-time detection and minimal sample preparation, nanopore sequencing is transforming genomics and proteomics [3, 5–7]. Its versatility has extended applications from clinical diagnostics and epigenetic profiling to field-deployable biosensing, marking it as a cornerstone of next-generation molecular analysis [8–10]. Over the past three decades, continual efforts have been devoted to engineering protein nanopores for improved performance and functionality [4, 11]. However, most rational designs remain limited to a few well-characterized protein nanopores, such as MspA and CsgG (Figure 1B), due to the technical difficulty and high cost of resolving membrane protein structures, particularly the narrow constriction zones essential for function [11, 12].

**Figure 1.**
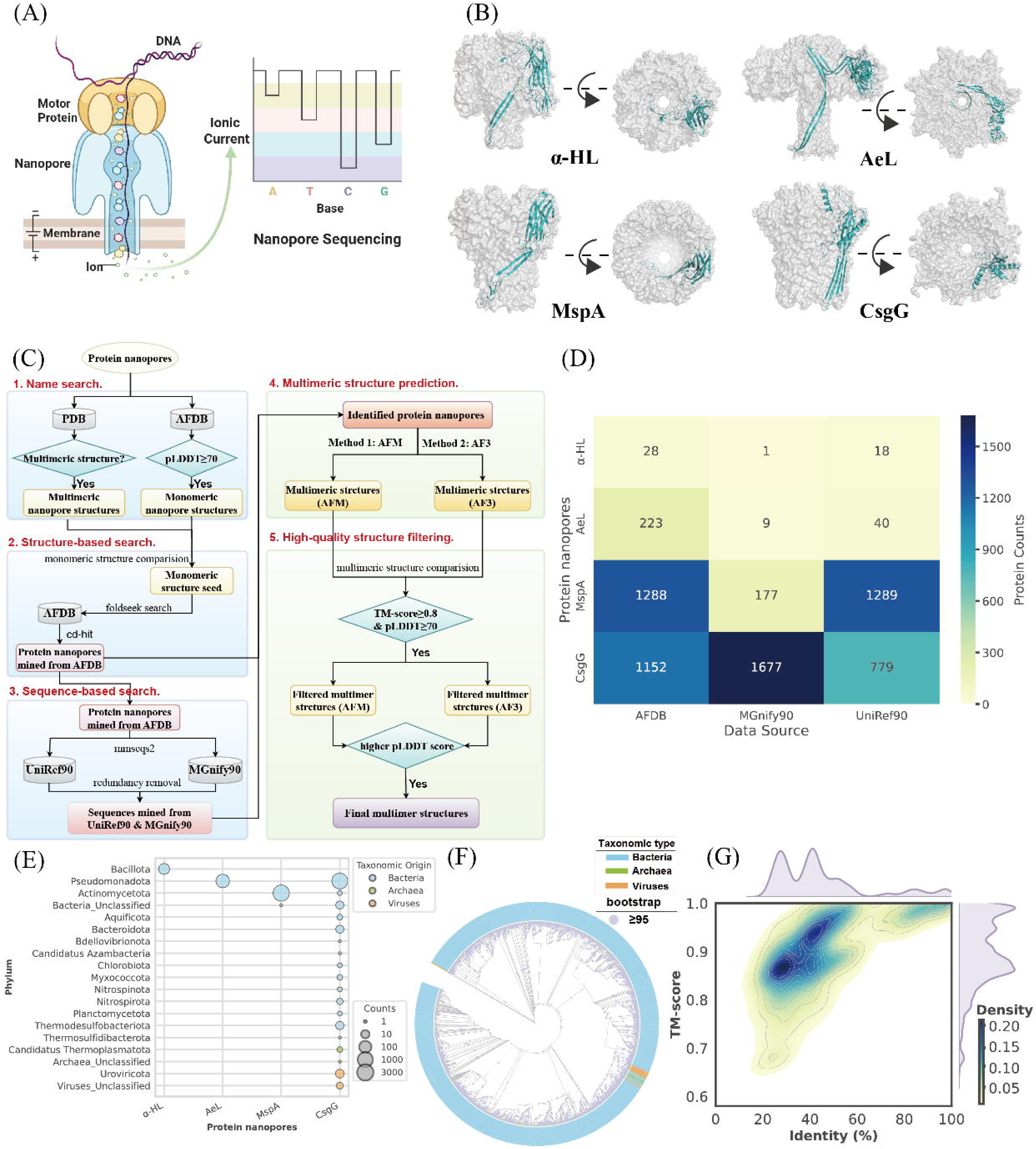
Structure- and sequence-based identification of protein nanopores. (A) Schematic illustration of nanopore structure and the principle of nanopore sequencing. (B) Structural representation of multimeric protein nanopores: α-HL (PDB: 7AHL), AeL (PDB: 9FM6), MspA (PDB: 1UUN), and CsgG (PDB: 4Q79). A monomeric chain of each nanopore is highlighted in cyan. (C) Overview of the nanopore mining workflow. (D) Source distribution of mined candidate protein nanopores. (E) Phylum-level distribution of annotated protein nanopores. The bubble chart displays protein nanopore counts across different phyla (y-axis) and nanopore types (x-axis). Bubble size corresponds to the number of proteins identified within each nanopore type. Blue, green, and orange bubbles indicate bacterial, archaeal, and viral phyla, respectively. (F) Phylogenetic tree of the identified CsgG proteins with taxonomic annotations. Ring colors represent the taxonomic origin of each sequence: blue for Bacteria, green for Archaea, and orange for Viruses. Purple circles on internal nodes denote branches with strong phylogenetic support (bootstrap ≥ 95). (G) Sequence-structure similarity landscape of candidate nanopores compared to representative PDB nanopore structures. The x-axis represents sequence identity, and the y-axis represents structural similarity (TM-score). Color gradient from light yellow to dark blue indicates increasing density of candidates.

The discovery and engineering of protein nanopores rely heavily on assessing structural features such as pore geometry and membrane embedment. Recent breakthroughs in deep learning-based methods for protein structure prediction, such as AlphaFold [13–15], enable fast acquisition of protein structures accurate enough for downstream rational design. In addition, large databases of predicted protein structures, such as AlphaFold Protein Structure Database (AFDB) [16] and ESM Metagenomic Atlas [17], offer a unique opportunity for mining new candidates of nanopores with improved sequencing capability. However, several issues of the current databases make them not readily usable for the discovery of novel protein nanopores. First, AFDB and ESM Metagenomic Atlas only provide predictions of single-chain proteins, whereas commercialized nanopores used for sequencing are large homomultimers, such as CsgG [18]. Because the predicted structures of monomers can alter after assembly, direct prediction of the multimeric structure is advantageous in inspecting the interior of the pore lumen. Second, derivative databases of AFDB, such as TmAlphaFold [19], MembranomeX [20], ChannelsDB 2.0 [21], AFTM [22], are focused on transmembrane alpha-helical proteins but contain few protein nanopores.

Here, to address the aforementioned limitations, we developed an open-access resource NanoporeDB, which is dedicated to systematically expanding the structurally annotated nanopore resource and provides a foundation for nanopore discovery and rational design. To construct NanoporeDB, we developed an integrated structure- and sequence-guided mining workflow that systematically identified homologous proteins from AFDB [16], UniRef90 [23], and MGnify90 [24], using experimental structures of nanopores from Protein Data Bank (PDB) [25] as references.

Candidate homologs were subjected to multimeric structure prediction using a locally-optimized AlphaFold-Multimer (AFM) [14] and AlphaFold3 (AF3) [15], and high-confidence structural models were retained for downstream analyses, including membrane embedding and pore geometry annotation. Compared to the few experimentally resolved multimeric structures currently available in the PDB (Table S1), NanoporeDB contributes over 6,600 high-quality predicted models, expanding the structurally annotated nanopore repertoire by more than 200-fold. We anticipate that NanoporeDB will serve as a valuable resource to advance the exploration and engineering of nanopores, enabling next-generation innovations in molecular sensing, precision diagnostics, and synthetic biology.

## Results

### 1. Workflow for protein nanopore mining

In this study, we developed a systematic mining workflow for a comprehensive encompass of protein nanopores of interest (Figure 1C). The pipeline consists of five consecutive steps: (1) Name search. We retrieved protein nanopores by name from both PDB and AFDB. (2) Structure-based search. We aligned the predicted monomeric structures from AFDB against the experimentally resolved templates from PDB and saved only the highly similar structures (methods). The retained multimer templates and predicted monomers were combined as structural seeds to align search against the entire AFDB for candidate structures (methods). (3) Sequence-based search. All unique sequences from step 2 were used to search against UniRef90 and MGnify90. (4) Multimeric structure prediction. The candidate sequences obtained from step 2 and 3 were merged and deduplicated for multimeric structure prediction by AFM and AF3, respectively. (5) High-quality structures filtration. All predicted models were aligned to the corresponding multimeric templates from PDB, whereas for each protein only the model with higher pLDDT score was retained to avoid redundancy (methods).

We selected four protein nanopores widely used in nanopore sequencing technologies on purpose, i.e., alpha-hemolysin (α-HL) [26], aerolysin (AeL) [27], MspA [28], and CsgG [29, 30]. Despite their significance, only a limited number of pore-like structures have been experimentally resolved for each protein type (Table S1), highlighting the urgent need for expanding the protein nanopore repertoire. Using our systematic mining workflow, we identified in total 6,681 candidate protein nanopores, including 47 α-HL, 272 AeL, 2,754 MspA, and 3,608 CsgG proteins (Figure 1D, Table S2), substantially broadening the available pool of protein nanopores for downstream structural and functional exploration. Among the nanopores with available taxonomic annotations from AFDB and UniRef90, the majority of α-HL, AeL, and MspA proteins were annotated to a single predominant phylum, reflecting a taxonomically concentrated distribution (Figure 1E). In contrast, CsgG-like nanopores displayed a broader taxonomic span, with the majority assigned to bacteria (98.8%) and a smaller number originating from archaea and viruses (Figure 1E). Notably, the archaeal and viral homologs are nested deeply within bacterial lineages (Figure 1F), indicating potential horizontal gene transfer of CsgG-like nanopores across distant evolutionary lineages.

To evaluate the diversity of the candidate nanopores, we compared their structural and sequence similarities to experimentally resolved multimeric protein nanopores from the PDB (Figure 1G, methods). Across all nanopore types, structural similarity consistently exceeded sequence similarity, indicating the conservation of nanopore structures despite considerable sequence divergence. These results demonstrate that our structure- and sequence-based mining workflow effectively detected remote homologs with conserved structural features but novel sequences, thereby expanding the protein nanopore pool beyond the reach of sequence-based strategies alone.

### 2. Assessment of predicted multimeric models

We assumed that predicting the multimers as a whole would improve the model quality of its constituent monomers compared to predicting the monomer alone when the inter-chain interactions were accounted for by AlphaFold. Therefore, we first compared the average pLDDT scores of the first chains of our predicted multimeric structures with their corresponding monomeric structures from AFDB (Figure 2A). In general, the multimeric models predicted by AFM and AF3 had monomers of better quality than their counterparts in AFDB for α-HL, MspA, and CsgG, indicating improved local structural confidence (Figure 2B). For α-HL, the median average pLDDT score was increased from 0.86 of the monomeric models from AFDB to 0.97 and 0.97 of the first chain of multimeric models generated by AFM and AF3, respectively. The difference in pLDDT between AFM models and AF3 models was insignificant. Similarly, MspA models showed improvement with the median pLDDT scores of 0.78, 0.93 and 0.91 for AFDB, AFM and AF3 models respectively, in which AFM slightly outperformed AF3. And CsgG nanopores exhibited a progressive increase in confidence, with median pLDDT scores of 0.81, 0.86, and 0.88 in AFDB, AFM, and AF3, respectively. Curiously, although AF3 generated multimer models of AeL with significantly better monomer than AFM, the corresponding monomeric structures in AFDB had the highest quality.

**Figure 2.**
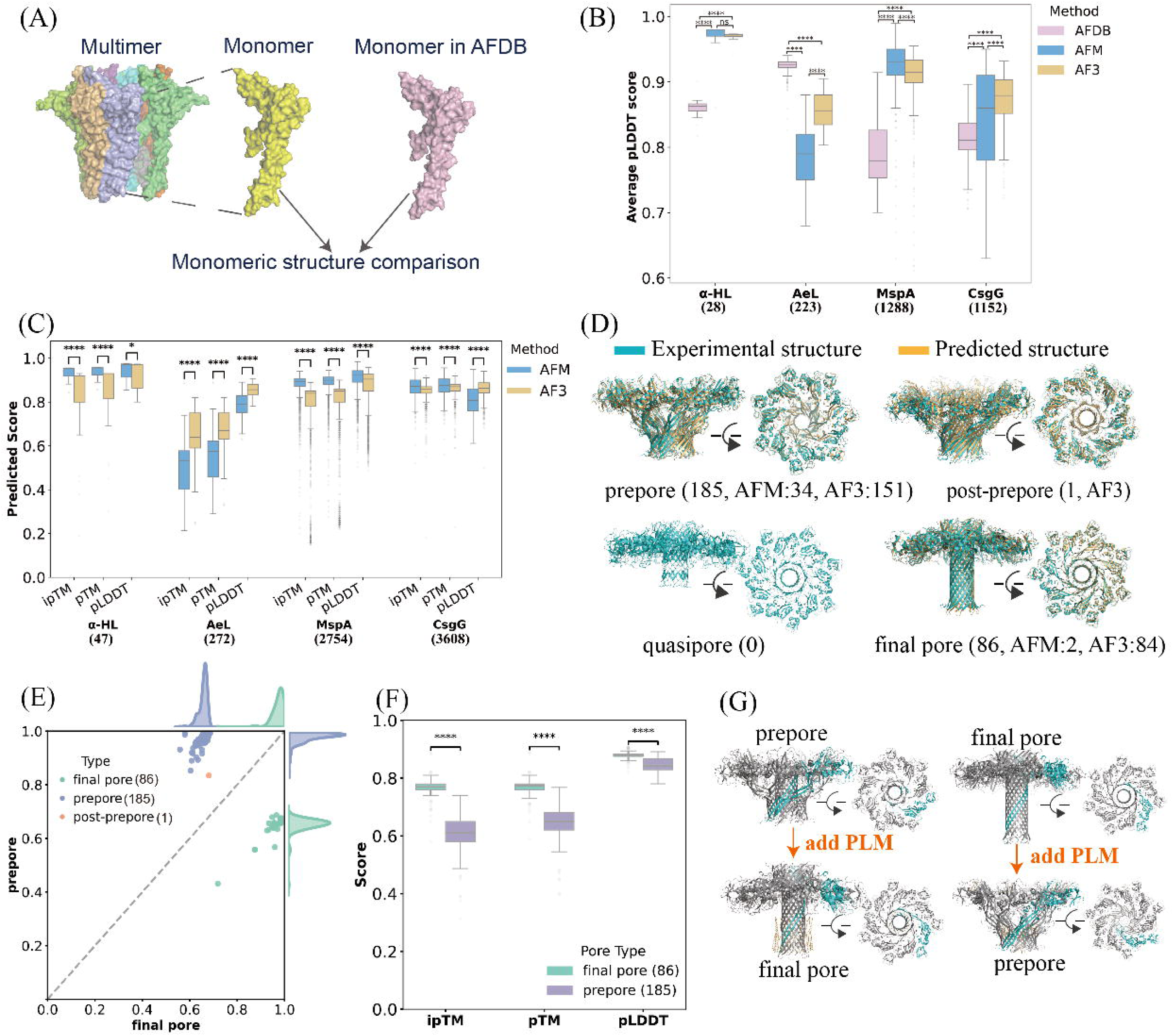
Evaluating predicted multimeric structures of identified protein nanopores. (A) Schematic of monomeric structure comparison. (B) Comparison of monomer-level average pLDDT scores across AFDB (purple), AFM (blue), and AF3 (yellow) models for each type of nanopore. Numbers in parenthesis indicate structures mined only from AFDB (Figure 1D). (C) Comparison of multimeric structure prediction scores (average pLDDT, pTM, and ipTM) between AFM (blue) and AF3 (yellow) models across all identified protein nanopores. Numbers in parenthesis indicate structures mined from all resources (Figure 1D). (D) Schematic representation of the four AeL conformations: prepore, post-prepore, quasipore and final pore. Reference structures (cyan) correspond to experimentally resolved states (prepore: 9FMX, post-prepore: 5JZW, quasipore: 5JZW, final pore: 9FML). Predicted AeL-like models with the highest similarity are shown in light orange and aligned with the experimental structures. Numbers in parenthesis indicate the models attributed to each conformational state and its predictor (Figure 1C). (E) Conformational grouping of AeL-like models. Scatter plot shows TM-scores of predicted models relative to a prepore (y-axis) and a final pore (x-axis). (F) Multimeric self-confidence scores (ipTM, pTM, and pLDDT) for the AeL prepore and final pore conformations. (G) The conformational transitions by adding PLM molecules (orange) in joint structure prediction. A monomer chain of each nanopore is highlighted in cyan. In B, C and F, significances of paired comparisons were calculated using two-sided Wilcoxon–Mann–Whitney U-test (\**P* < 0.05, \*\*\*\**P* < 0.0001).

We then assessed the quality of the overall multimer models predicted by AFM and AF3 by comparing three structural metrics: ipTM measures the quality of interfaces between interacting monomers, pTM evaluates the general fold of the multimeric structures, and the average pLDDT scores indicate the confidence of local structures (Figure 2C). In parallel to the above monomer comparisons, for α-HL, MspA and CsgG nanopores, both methods yielded consistently high scores (> 0.8) across all three metrics (Figure 2C). In contrast, AeL nanopores exhibited markedly lower scores across all three metrics compared with the other pore types, and within this group AF3 significantly outperformed AFM (Figure 2C).

The relatively lower confidence scores observed for AeL nanopores (Figure 2B, C) prompted us to examine the potential impact of the conformational heterogeneity. During pore formation, the AeL proteins undergo a series of conformational changes: from prepore to final pore via two transient intermediate states, i.e. post-prepore and quasipore (Figure 2D, Table S1) [31]. This progress begins with the oligomerization and formation of two stable concentric β-barrels (prepore); then, the protein morphs though a zipper-like formation and piston-like extension of the inner β-barrel (post-prepore and quasipore), and finally inserts into the lipid bilayer (final pore) [31]. Our workflow predominantly captured the prepore (185 of 272 models, 68%) and final pore (86 of 272 models, 32%) states based on the global structural similarity, whereas the post-prepore (1 model) and quasipore (0 model) states were underrepresented (Figure 2D, E, methods). Further, we showed that the models adopting the final pore conformation consistently achieved high self-confidence scores, indicating that the lower overall confidence of AeL models primarily resulted from the dominance of prepore conformations, which were intrinsically more flexible and heterogeneous prior to membrane insertion (Figure 2F, Figure S1).

Given the scarcity of these intermediate states in prediction, the subsequent analyses were focused on comparing the prepore and final pore states. Because our workflow would select the multimer model with higher self-confidence from the AFM and AF3 predictions (Figure 1C), we noticed that both the prepore and the final pore conformations were represented mostly by AF3 models (Figure 2D). We hence compared the two predictors. All AeL models predicted by either AFM or AF3 were aligned against the reference experimental structures of the prepore and final pore, respectively (Figure S2, Table S3, methods). AFM models were strongly biased toward the prepore state, with about 80% exceeding a TM-score of 0.7 to the reference prepore structure and only a negligible fraction matching the reference final pore structures (Figure S2A, Table S3). In contrast, AF3 predictions were distributed across both states, with 67% inclined to the prepore and 31% to the final pores (Figure S2B, Table S3), underscoring AF3’s capability of broader conformational sampling.

We further examined whether the addition of membrane-mimicking components could alter the conformational switching (Figure 2G). Hence, we introduced palmitic acid (PLM) as a lipid surrogate in AF3 predictions, which has a polar head and the longest lipophilic fatty tail available on AF3 online server. In total, 46 prepore-like AF3 models transitioned to the final pore state in the joint prediction of nanopore plus PLM in which cases PLM molecules formed a bilayer, whereas 4 final pore-like models reverted to the prepore state with slightly reduced self-confidence (Figure 2G, Figure S3A, Table S4). Though predictions with PLM showed elevated pLDDT values, adding PLM did not improve ipTM and pTM scores (Figure S3B), indicating that introducing additional entities in AF3 prediction affected differently the local and global structure. Further, the observation that adding lipophilic entities in joint structure prediction biased protein conformation towards the transmembrane state (i.e. the final pore state) naturally led us to postulate the opposite scenario of adding hydrophilic entities. Thus, we replaced PLM molecules with potassium and chloride ions (Figure S4). As expected, we observed 84 models with conformational changing: 64 from the final pore state to the prepore state, and 4 vice versa (Table S4).

### 3. Protein nanopore structural analysis

To facilitate the application of candidate protein nanopores in single-molecular sequencing, we performed in-depth structural analysis, including membrane embedding and pore constriction analysis (Figure 3A, B). The latter was focused on computing pore radius and the axial location along the pore channel (Figure 3B). We first computed the insertion and orientation of each multimeric complex within a model lipid bilayer (1,2-dioleoyl-sn-glycero-3-phosphocholine, i.e. DOPC) using PPM 3.0 [32]. For each model, we computed its key structural parameters including the insertion depth and the tilt angle of the pore axis relative to the membrane normal. It is noted that we only included AeL-like candidates exhibiting the final pore conformations in membrane embedding analysis (Figure 2D), because the prepore state should not entered the membrane [31]. As nearly 68% of AeL-like models were in prepore-like, the conformation-based filtering was essential to evaluate nanopore candidates. Nonetheless, being in the final pore state did not guarantee membrane insertion (Figure S5B, S6A). It cautions us to rely on multiple criteria in candidate assessment.

**Figure 3.**
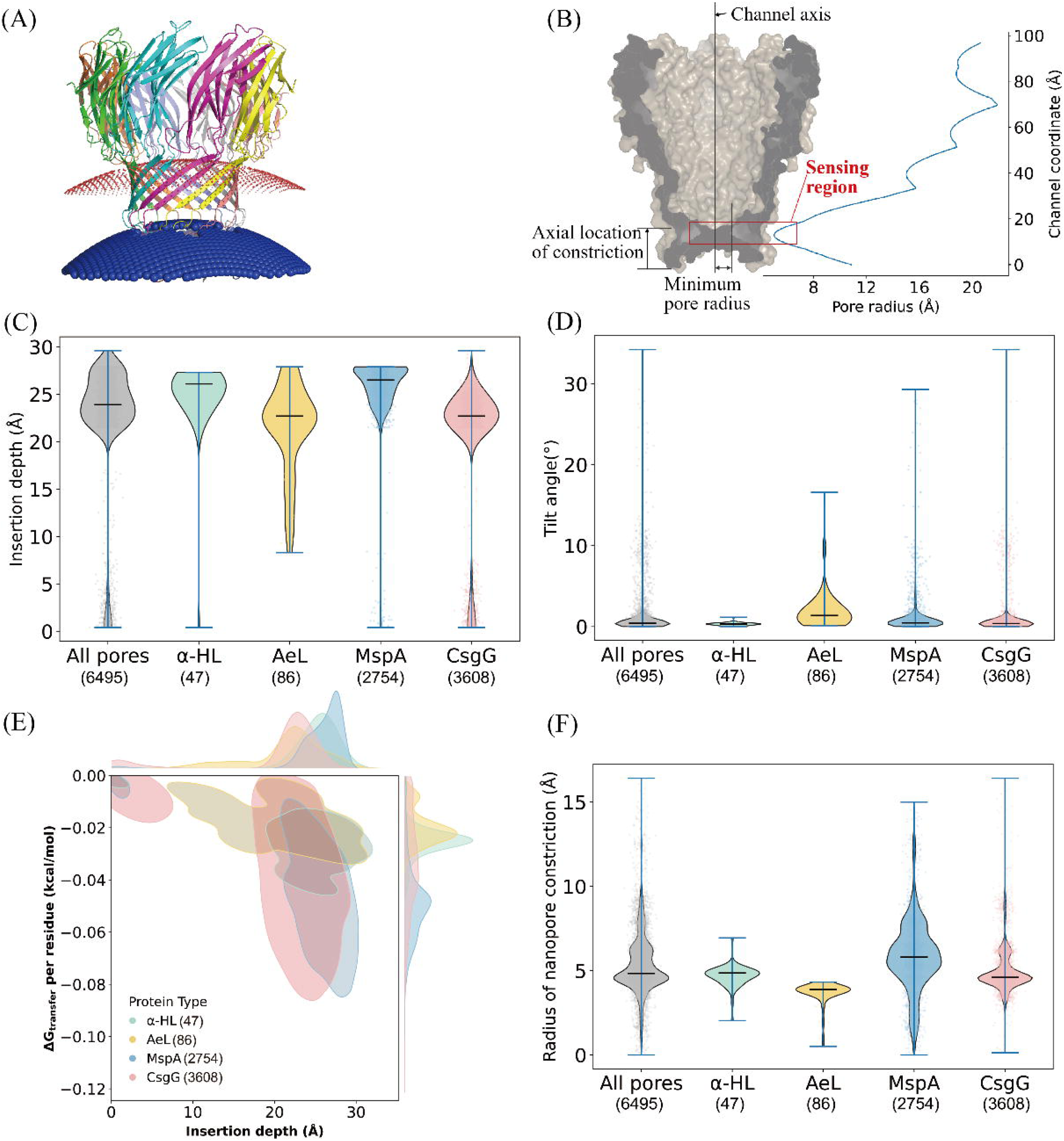
Membrane embedding and structural features analysis of candidate protein nanopores. (A) Example of predicted nanopore embedding. Red and blue dots stand for the outer and inner membranes, respectively. (B) Example of the minimum pore radius and its corresponding axial location along the nanopore channel. Representative profile of pore radius along the channel axis for an MspA-like nanopore (AFDB ID: A0A064C8X8). (C) Violin plot of predicted insertion depth (Å) for nanopore candidates. (D) Violin plot of predicted tilt angles (°) relative to the membrane normal (methods). (E) Density plot of transfer free energy per residue for nanopore candidates. (F) Violin plot of predicted minimum pore radius values (Å) across nanopore candidates.

Across all analyzed complexes, the predicted nanopores were embedded in lipid bilayers with a median insertion depth of 23.9 Å (Figure 3C). The central pore axes were uniformly oriented perpendicular to the local membrane surface with a median tilt angle = 0.4° (Figure 3D) with a few exceptions (Figure S6). Nanopore-specific variations were observed: α-HL and MspA-like pores showed deeper insertions (median ≈ 26 Å), whereas AeL- and CsgG-like pores exhibited shallower embeddings (median ≈ 23 Å), reflecting pore-specific hydrophobic matching. A minority of all models (11.0%) displayed anomalous shallow insertion depths (< 20 Å) (Figure 3C, Figure S5). For MspA-like pores, such cases likely reflect possible multiple conformations similar to that of AeL (Figure S5C); whereas for CsgG-like pores, those cases may result additionally from the absence of the native N-terminal membrane anchor (Figure S5D), which was previously shown to be essential for bilayer insertion [29]. From the perspective of energetics, the nanopore-specific differences were shown in the distribution of transfer free energy per residue: pores with deeper insertion generally had more favorable insertion energies (Figure 3E). MspA exhibited the greatest mean insertion depth (Figure S7), which more closely matched the hydrophobic thickness of DOPC bilayer [33], and thereby minimized the hydrophobic mismatch to obtain the most favorable transfer energy (Figure 3E).

To further gauge the structural suitability of the candidate protein nanopores for potential analyte translocation, we analyzed their pore geometries. We calculated the pore radius along the central channel axis to identify the location and width of the narrowest constriction zone (Figure 3B, F), which are critical determinants for sensing resolution and analyte compatibility [34]. The predicted MspA-like model displayed consistent features with the canonical MspA nanopores [35], including a minimal pore radius of approximately 5 Å positioned at ∼13 Å above the β-barrel terminus along the channel axis (Figure 3B). Narrow constriction is a known characteristic to enhance single-molecule signal discrimination during translocation [28, 36]. In general, the predicted minimum pore radii exhibited a wide range (0.2 Å to 16 Å), with the vast majority (> 93%) of structures falling within the 3 to 10 Å range (Figure 3F). This size range permits the translocation of single-stranded DNA and small peptides [28, 37], suggesting the potential for further engineering toward single-molecule sensing applications. While the majority of candidates held promise for sensing applications akin to current nanopore technologies, the existence of pores with both smaller and larger constrictions hinted at a wider functional potential that merits further investigation (Figure S8).

### 4. NanoporeDB: a web-based resource for structural profiling of protein nanopores

To facilitate the discovery and structural evaluation of protein nanopore candidates, we developed an interactive web platform NanoporeDB (Figure 4A). For each entry, NanoporeDB provides the multimeric assembly predicted by either AFM or AF3, which can be freely downloaded on the entry page (Figure 4B). Structural features relevant to nanopore sensing, including the minimum pore radius and its corresponding axial position relative to the β-barrel terminus, insertion depth, tilt angle, and transfer free energy, are displayed as well (Figure 4B). Examining these parameters (e.g., minimum pore radius < 6 Å, tilt angle < 10°) can allow users to readily identify nanopore candidates that meet the dimensional requirements for the intended sensing or sequencing applications.

**Figure 4.**
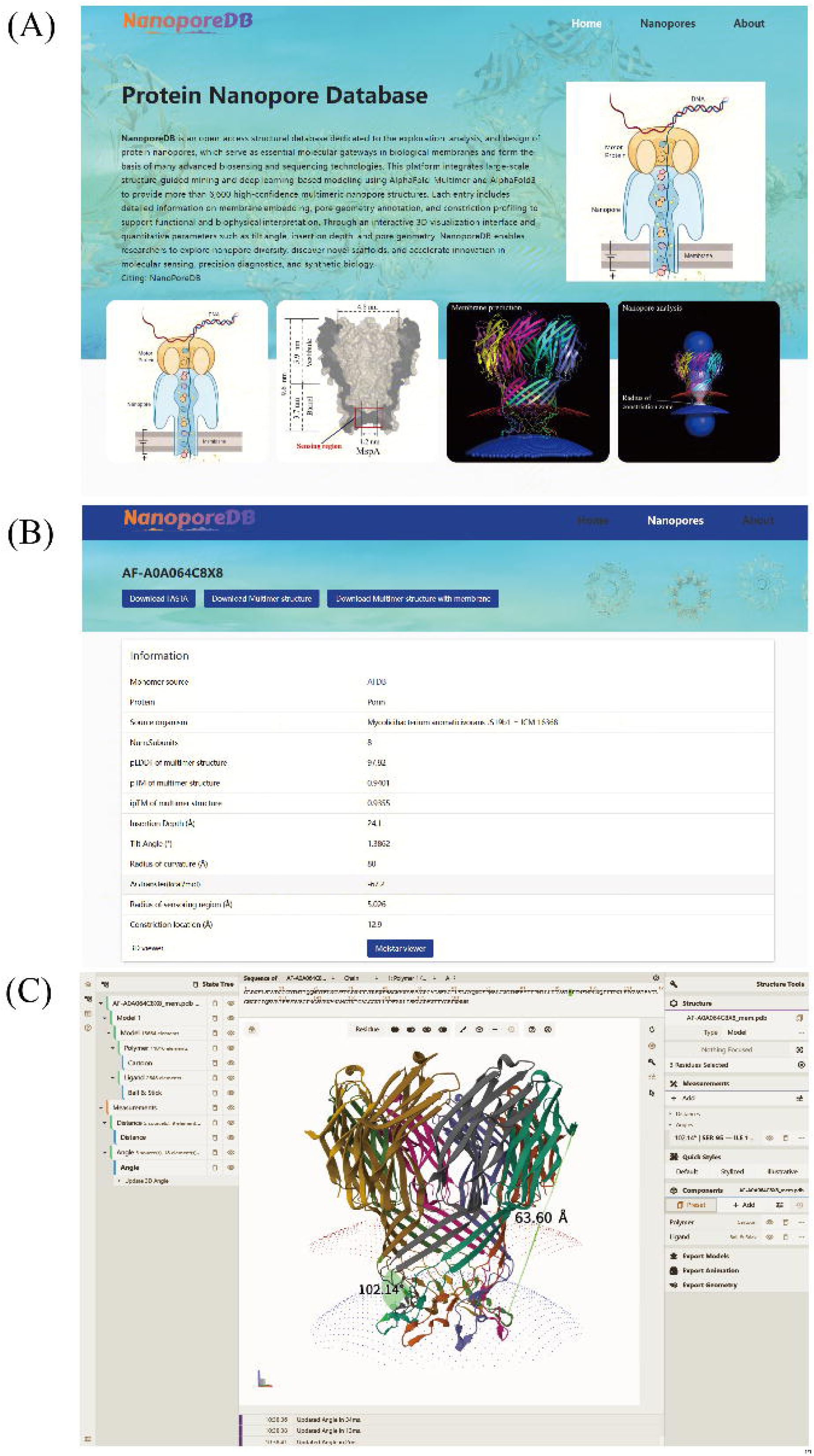
Web interface and data visualization of NanoporeDB. (A) Homepage of the NanoporeDB website. (B) The webpage of a deposited nanopore candidate detailing the information for individual protein nanopores, including sequence, structure, and structure annotations. (C) Example of the 3D visualization UI of a membrane-embedded protein nanopore structure.

In addition, we implemented an interactive “3D viewer” user interface (UI) enabling flexible online inspection of pore structure for each entry (Figure 4C). Users can conveniently explore the multimeric nanopore structures at multiple levels, from individual atoms and residues to complete chains. Beyond flexible selection and visualization, the UI supports precise structural measurements such as the distances between residue pairs, bond lengths and dihedral angles among selected residues (Figure 4C). The integrated visualization, along with downloadable model files and detailed structural metrics, enables efficient screening and rational design of nanopore-based biosensing and sequencing applications.

## Discussion

Traditional sequence-based mining approaches are often limited in detecting remote homologs, as evolutionary divergence rapidly erodes sequence similarity while leaving structural features comparatively conserved. Recent studies have demonstrated that structure-based or structure-informed strategies can substantially expand the discovery of novel protein families, such as novel deaminases identified through structure clustering [38] and TIGR-Tas systems uncovered by integrating predicted structures with sequence features [39]. Consistent with these observations, our results show that structural similarity among nanopore candidates was consistently higher than sequence similarity (Figure 1G), demonstrating the effectiveness of structure-guided mining in identifying novel protein nanopores and extending the protein nanopores beyond the reach of sequence-based approaches.

Among the nanopore candidates, α-HL and AeL-like nanopores are largely associated with pathogenic lineages, and MspA is primarily confined to the Actinomycetota (Figure 1E). In contrast, CsgG-like proteins exhibited a markedly broader taxonomic distribution compared to α-HL, AeL, and MspA nanopores, with homologs detected across diverse bacterial phyla, as well as in archaea and viruses. This widespread occurrence likely reflects the conserved role of CsgG as the outer-membrane channel of the curli secretion system, which is essential for biofilm formation and environmental adaptation in many Gram-negative bacteria [40, 41]. The widespread distribution of CsgG homologs suggests possible horizontal gene transfer, as secretion-system components are known to be prone to horizontal transfer [42]. For instance, metagenomic surveys of seawater phages showed strong enrichment of adhesion-related genes with CsgG being the most abundant [43].

Our comparative analysis revealed that for protein nanopores the monomeric subunits extracted from AFM- and AF3-predicted multimeric structures generally exhibited higher average pLDDT scores than their monomeric counterparts from the AFDB (Figure 2B). This improvement in local structure prediction demonstrates that the explicit incorporation of inter-chain interactions during multimeric folding can refine model quality of the constituent monomers, which may result from resolving conformational ambiguities at multimeric interfaces and within pore-lining regions that remain poorly captured in isolated monomer predictions. Admittedly, more reliable interface-specific metrics may offer better comparisons of predicted complex models [44–48]. Consistent observations across α-HL, MspA, and CsgG-like nanopores emphasize the necessity of multimeric modeling for protein nanopores (Figure 2B). Moreover, modeling the entire multimer offers the particular advantage of illuminating regions that are often unresolved in experimental structures. For example, the secretin GspD of the type II secretion system is a huge pentadecameric nanopore [49] and has been reported as a competent next-generation candidate for DNA sequencing applications [50, 51]. Multimer structure prediction can provide valuable hypothesis [52] of the unresolved key linkers in the central and cap gates of GspD which are critical for its sensing ability [49].

On the other hand, AeL-like proteins represented an informative exception that merits further consideration (Figure 2). First, AFM predictions were strongly biased toward the prepore state (Figure S2A), whereas AF3 models were distributed in both conformational states in a balanced manner (Figure S2B), indicating AF3’s advantageous performance in conformational sampling for certain proteins with multiple intermediate states. But AF3 is known for its limited ability in distinguishing conformational switching [15, 53]. Besides, neither approach yielded better monomer in complex prediction in either prepore or final pore state than the monomeric model from AFDB which was directly predicted by AF2 (Figure S1A). These reflect the intrinsic challenge in predicting the multimeric structure of protein nanopore, especially when it undergoes large conformational transitions during the embedment into membrane [31]. The long-tailed distributions in ipTM and pTM scores of MspA and CsgG (Figure 2C) likely suggest existence of prepore intermediates yet to be captured by experiment (Figure S5C left, S5D left). Second, incorporation of membrane-mimicking components in AF3 effectively modulated the outcomes: PLM promoted conformational shifts toward final pores and hence rescued membrane embedding that 65 out of 127 (51.1%) AeL-like final pores were able to be inserted into bilayer correctly (Table S4), among which there were 23 prepores switching to the final pore state, while there were only 20 out of 84 (23.8%) final pores were able to embed correctly (Table S4). Similar co-folding outcomes were seen in 26 out of 32 MspA and 19 out of 41 CsgG models (Figure S9, Table S2, methods). Still, the co-folding strategy does not promise protein models of better quality in general (Figure S3B, S4). Though previous works demonstrated that AF3 did not learn the physics but memorized ligand-protein interactions instead [53–55], our observations suggest that environmental surrogates can sway conformational sampling and hence produce desirable models. In general, accurately predicting multiple conformations of assemblies with morphing domains requires physics-informed structure prediction methods, possibly in combination with emulation of protein equilibrium ensembles [56].

Together, these findings establish NanoporeDB as a valuable resource for accelerating nanopore discovery and design in biosensing, sequencing, and diagnostic applications. Nevertheless, several limitations of our current version should be acknowledged. First, although AlphaFold-based models of protein nanopores generally exhibit high quality, experimental validation is essential to confirm their ability to capture functional conformations and properties relevant to desired applications. Second, the structural analysis of pore geometry depends critically on robust determination of the channel axis, which may be affected by local asymmetry or structural flexibility in some multimeric assemblies. Thus, manual curation of the pore axis served as a complement to automatic computing tools, such as HOLE [57], to ensure reliable radius profiling across the diverse nanopore structures. Third, membrane embedding was modeled under simplified conditions, assuming a static protein conformation and homogeneous lipid bilayer. In reality, biological membranes are compositionally heterogeneous and dynamically fluid, with lipid–protein interactions lateral pressure variations, and local curvature, which potentially affect pore orientation, stability, and function [58, 59]. Forth, our multimer prediction presumed the same stoichiometry as the experimental structure of given protein nanopore. However, isoforms of transmembrane channels are known to form diverse oligomeric assemblies [60]. Thus, some of our predicted models with small pore radii may in fact assemble with more monomers, or vice versa. Fifth, the current version of NanoporeDB merely encompasses a limited set of nanopores, primarily focusing on representative nanopore types with commercial applications. The future release of NanoporeDB will expand the scope to include additional pore types such as GspD [50, 51].

## Conclusion

In summary, we established an integrated workflow for the discovery and structural profiling of four representative types of protein nanopores, and developed NanoporeDB as the first online resource to provide membrane-embedded multimer models for over 6,600 protein nanopores with annotated pore geometries. By democratizing access to these structurally annotated multimeric nanopore models, NanoporeDB will directly accelerate innovation in sequencing technologies and impact fields reliant on these advances, including clinical diagnostics, infectious disease surveillance, epigenetic research, single-molecule proteomics, etc.

## Methods

### Data collection

To identify protein nanopores on a large scale, we collected data from several high-quality structural and sequence databases. Experimentally determined protein structures were retrieved from the PDB (available as of March, 2025) [25], while predicted protein structures were obtained from the AFDB v4 (available as of March, 2024) [16]. Only AFDB monomeric models with high confidence (average pLDDT ≥ 70) were considered. In addition to structural data, we used UniRef90 (version: November, 2024) [23] and full-length protein sequences from MGnify90 (version: April, 2024) [24] as comprehensive protein sequence databases for downstream homology searches.

### Protein Nanopore Mining workflow

To systematically identify and model protein nanopores from large-scale structure and sequence databases, we established a five-step computational workflow (Figure 1C).

1. Name-based retrieval. Representative nanopore proteins, including α -hemolysin (α-HL), aerolysin (AeL), MspA, and CsgG, were first retrieved by name from the PDB and AFDB v4 [16]. Only experimentally resolved multimeric pore-forming assemblies were retained to ensure structural relevance. These served as initial structural templates for subsequent searches.
2. Structure-based search. Predicted monomeric structures from AFDB were aligned to each experimentally resolved nanopore monomer from PDB using the *easy-search* module of Foldseek (v9.427df8a) [61]. Only hits with TM-score ≥ 0.8 were retained to ensure high structural confidence. The retained AFDB monomers and PDB templates were then merged as structural seeds for an expanded search across the entire AFDB database, enabling the identification of potential homologous nanopores with conserved structures but divergent sequences.
3. Sequence-based search. All unique sequences obtained from the structure-based search were further used as queries against UniRef90 and MGnify90 databases using MMseqs2 (commit: 0b27c9d7d7757f9530f2efab14d246d268849925) [62] (-c 0.9 -e 1e-4). This step expanded the nanopore candidates by identifying distant homologs sharing sequence or structural similarity.
4. Multimeric structure prediction. Candidate sequences from steps 2 and 3 were merged and deduplicated. Multimeric structures were predicted using AlphaFold-Multimer (AFM, locally optimized implementation) [14] and AlphaFold3 web server (released March 18, 2025) [15]. Predictions were performed under identical subunit stoichiometries as the corresponding experimental templates.
5. Quality filtering and redundancy removal. All predicted multimeric structures were aligned to their corresponding PDB templates using US-align (v20241108) [63]. Only models with TM-score ≥ 0.8 were retained. For cases with multiple predictions of the same sequence, the structure with the highest pLDDT score was selected as the representative nanopore model.

### Monomeric Structure Comparison

To assess the structural similarity of predicted monomeric units, we used Foldseek *easy-search* (-c 0.7 -e 0.01 --tmscore-threshold 0.8 --alignment-type 1 --format-output query,target,alntmscore,qtmscore,ttmscore,qcov,tcov,lddt,prob). For each pairwise comparison, monomeric models with either qtmscore or ttmscore > 0.8 were retained as high-confidence structural matches.

### Multimer structure prediction

To predict the multimeric structures of candidate protein nanopores identified through sequence- and structure-based mining, we employed both AFM and AF3. Prior to structure prediction, we applied a truncation strategy to improve prediction accuracy and reduce computational load. Specifically, each candidate sequence was aligned to structurally resolved protein nanopores in the PDB using BLAST+ 2.16.0 [64], and the closest homolog (with the highest sequence identity) was selected as the reference. Residues extending beyond the N-and C-terminal boundaries of the reference structure were removed to eliminate potentially disordered or irrelevant regions that could interfere with accurate structure prediction. The refined sequences were then subject to AFM and AF3 for multimeric structure prediction. Both methods provided accurate and efficient structural prediction across different types of nanopores with diverse oligomeric architectures. We noted that longer sequences required substantially greater computational time and memory, highlighting the need for efficient sequence preprocessing prior to inference.

AFM was implemented as a GPU memory-optimized variant of AlphaFold-Multimer, capable of handling input sequences up to approximately 8,200 amino acids. Instead of directly modifying the original JAX-based AFM, which provides limited flexibility for GPU memory management, we developed our implementation based on Uni-Fold v2.2.0 [65]. Uni-Fold is an open-source PyTorch re-implementation of AlphaFold2 that maintains full compatibility with the official model parameters and supports both training and inference. It introduces a two-dimensional blocking strategy within the Triangular Multiplication module to reduce peak GPU memory usage, providing a more flexible and extensible foundation for large-scale structure prediction. Using the converted parameters *multimer_model_1* (released December 6, 2022) [14], structural inference was performed on NVIDIA A100 GPUs (80 GB). To overcome the default memory limitations of Uni-Fold, which typically fails for sequences longer than approximately 4,300 residues, we implemented a targeted memory optimization strategy. Redundant intermediate tensors were discarded, temporarily unused variables were offloaded to CPU memory and reloaded dynamically when required, and inference was conducted in bfloat16 precision to further reduce memory overhead. These enhancements collectively enabled stable prediction of multimeric complexes up to ∼8,200 residues, thereby facilitating the accurate modeling of large assemblies such as protein nanopores. The inference configuration used in this study was: *model_name* = multimer_af2_v3, *base_seed* = 42, *num_ensembles* = 1, *max_recycling_iters* = 3, *bf16* = True.

In parallel, multimeric structures were also predicted using AF3 via the official AlphaFold3 Server (https://alphafoldserver.com/, released March 18, 2025) [15]. As with AFM, input sequences were first truncated based on their closest structural homologs in the PDB. The number of protomers in each assembly was determined based on the canonical oligomeric state of the corresponding nanopore type (for example, heptamer for α-HL and AeL, octamer for MspA, nonamer for CsgG). Each processed sequence of the candidate was submitted to the AF3 server, which generated five predicted models. The model with the highest *ranking_score* was selected for downstream analyses. This strategy ensured that only the most reliable multimeric assemblies were retained for subsequent structural comparisons, pore annotation, and functional interpretation.

### Multimeric Structure Comparison

To assess the global similarity of predicted multimeric assemblies, we used US-align [63] (-mol prot -mm 1 -ter 1 -outfmt 2) to align each predicted multimeric structure against experimentally resolved multimeric structures from the PDB. Models exhibiting either *qtmscore* or *ttmscore* > 0.8 were retained, ensuring that only high-confidence matches with consistent quaternary topologies were considered. The retained multimeric structures were further analyzed for membrane embedding and pore geometry.

### Conformation assignment of AeL-like models

We compared each predicted multimeric model with all experimentally resolved structures (Table S1, S3) using US-align [63] (-mol prot -mm 1 - ter 1 -outfmt 2). Each model was assigned to the conformation yielding the highest *qtmscore* if *qtmscore* >= 0.7. If a model could not be aligned to any of the experimental structure, its conformational state was assigned as “other”, indicating a potential novel intermediate state.

### Phylogenetic Analysis of Identified CsgG Proteins

Protein sequences of identified CsgG nanopores with available taxonomic annotations (Bacteria, Archaea, and Viruses) were aligned using Clustal Omega (v1.2.4) [66] with default parameters. To root the phylogenetic tree, one homologous protein sequence (UniProtKB: Q1RD49) was included as an outgroup. The multiple sequence alignment was refined by removing poorly aligned positions and gaps using trimAl (v1.5) [67] with the -automated1 setting, producing a high-quality alignment. Phylogenetic reconstruction was performed with IQ-TREE 3 (v3.0.1) (-m MFP -B 1000 -alrt 1000). The resulting Newick tree was visualized and annotated in iTOL (v7.2.2) [68].

### Multimer structure prediction with PLM and ionic environments

Multimeric nanopore structures were predicted using AF3 in protein–ligand–multimer mode with ∼40 palmitic acid (PLM) molecules per model to mimic lipid surroundings. Control runs with 10 KL and 10 ClL ions to assess environmental effects on assembly formation. Model confidence was evaluated using pLDDT, pTM, and ipTM scores.

### Membrane embedding prediction

Membrane insertion prediction was performed for all high-confidence multimeric nanopore structures using PPM 3.0 [32]. We selected 1,2-dioleoyl-sn-glycero-3-phosphocholine (i.e., DOPC) as the reference lipid bilayer, which closely approximates the physicochemical properties of membranes typically used in nanopore sequencing experiments. PPM automatically determines the energetically favorable membrane orientation by minimizing the transfer free energy from aqueous to membrane environments. For each structure, PPM selected either a planar or curved bilayer model, depending on the structural curvature and energy landscape. The output included detailed structural parameters such as insertion depth, tilt angle, transfer free energy. These parameters were used to assess compatibility with lipid bilayers and downstream comparative analyses.

### Membrane embedding evaluation

To assess the orientation and membrane compatibility of predicted nanopores, we evaluated their membrane embedding geometry using the PPM 3.0 predicted models. For each multimeric structure, the nanopore channel axis was defined as described in the “Pore radius calculation”. The planar (or curved) bilayer surface generated by PPM 3.0 was then used to derive its local normal vector. The tilt angle was calculated as the acute angle between the nanopore channel axis and the membrane normal, representing the inclination of the pore relative to the membrane plane. A nanopore was defined as *“able to embed”* if it had residues located on both sides of the membrane bilayer, indicating successful transmembrane insertion. Among these, structures showing a tilt angle of ≤ 10° were further classified as *“able to embed correctly”*, reflecting near-perpendicular alignment to the bilayer. These metrics allowed us to access membrane compatibility and inform downstream analyses of pore geometry and structural comparability.

### Pore radius calculation

To characterize the internal pore architecture, we employed HOLE (v2016.8.7) [57] to compute the radius profile along each nanopore’s channel axis. Given that HOLE’s automated pore axis detection can produce suboptimal results for asymmetric or irregularly shaped pores, we defined the pore axis manually for each structure. Specifically, the β-barrel region was identified based on secondary structure annotations generated using the DSSP algorithm [69] as implemented in MDTraj (v1.10.3) [70], selecting continuous β-strand segments containing at least 10 consecutive residues. For each homomeric nanopore, we then calculated the geometric center of the entire structure as well as the centroid of the identified β-barrel region. The vector connecting these two points was used to define the axial direction, with the β-barrel centroid designated as the starting point of the pore scan.

This axis and origin were then provided as input to HOLE, which performed a scan in 0.1 Å steps along the defined axis until reaching the extracellular opening. To ensure robustness and mitigate sensitivity to termination thresholds, we repeated the analysis across a range of end distances (10–60 Å, in 2 Å intervals). We then selected the most consistent local minimum radius as the representative constriction point for each nanopore structure. This approach enabled reliable identification of the narrowest region within the channel, facilitating subsequent comparative and functional analyses.

## DATA AVAILABILITY

Website of NanoporeDB will be public upon publication of the paper.

## AUTHOR CONTRIBUTIONS

Conceptualization, Z. W., Yuqian L.; Data curation, Yuqian L., Z. S., J. Z., Z. W.; Formal Analysis, Yuqian L., Z. W.; Website construction, W. Y., Yuqian L., D. L., K. M.; Writing – Original Draft, Yuqian L., Z. W.; Writing – Review & Editing, all authors; Visualization, Yuqian L., Z. W., W. Y.; Project Administration, Z. W., Yong Z., T. Z.; Supervision, Y. D., X. X.; Funding Acquisition, X. X., Y. D., Yuning Z.

## ACKNOWLEDGEMENT

This work was supported by National Key R&D Program of China (No.2024YFC3406300), “Pioneer” and “Leading Goose” R&D Program of Zhejiang (2024C03004), and Shenzhen Science and Technology Program (KQTD20221101093603011).

## DECLARATION OF GENERATIVE AI AND AI-ASSISTED TECHNOLOGIES IN WRITING

During the preparation of this manuscript, the authors used Deepseek to polish the manuscript. After using this tool, the authors have reviewed and edited the content as needed and take full responsibility for the content of the publication.

