## Supplemental materials for "NanoporeDB: A Structural Resource Of Multimeric Protein Nanopores In Single-Molecule Sensing"

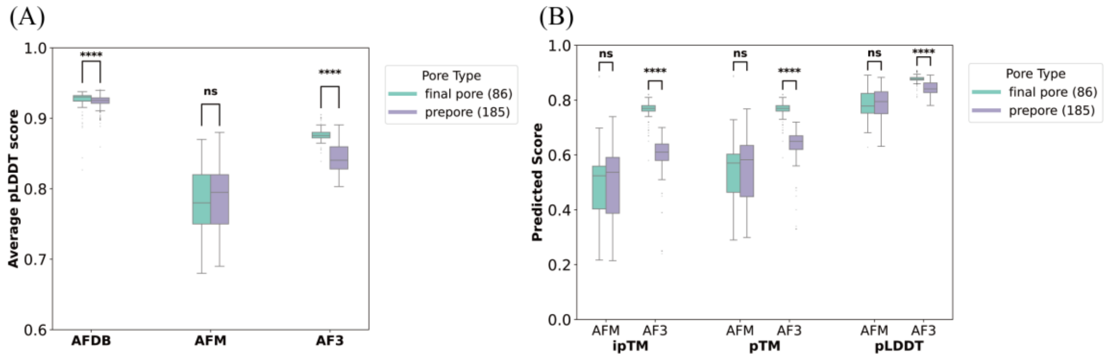


**Figure S1.** Comparison of self-confidence scores between AeL prepore (purple) and final pore (teal) models. (A) Monomer-level average pLDDT scores for AeL models from the AFDB, as well as those predicted by AFM and AF3. (B) Multimer-level scores (ipTM, pTM, and average pLDDT) of AFM and AF3 predictions. Significances of paired comparisons were calculated using two-sided Wilcoxon–Mann–Whitney U-test (*****P* < 0.0001).

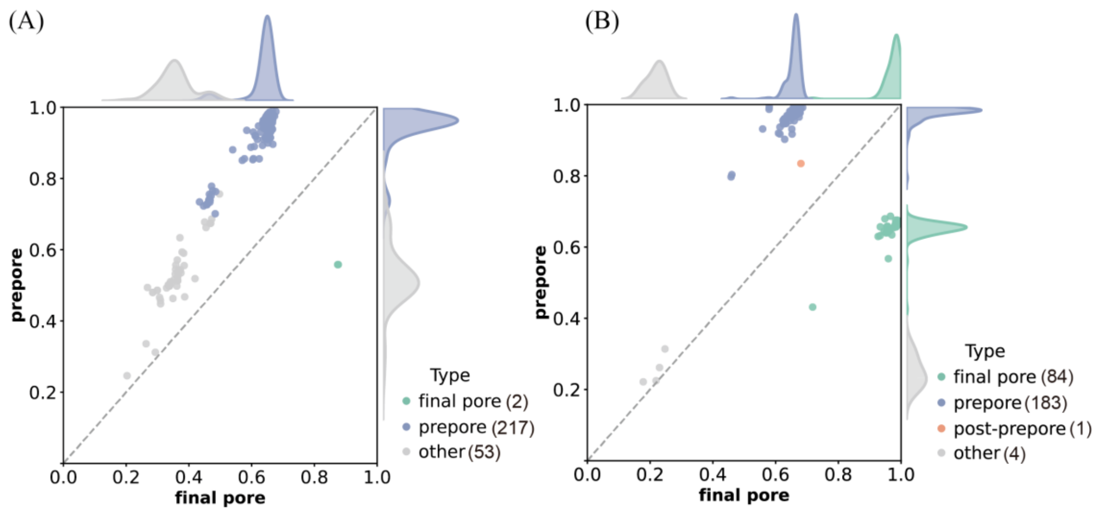


**Figure S2.** Conformational grouping of AeL-like models predicted by (A) AFM and (B) AF3**.** Scatter plots show TM-scores of predicted models relative to reference prepore (y-axis) and final pore (x-axis) structures. Points are colored by the assigned states. Models with TM-scores < 0.7 against the reference structures were classified as “other”.

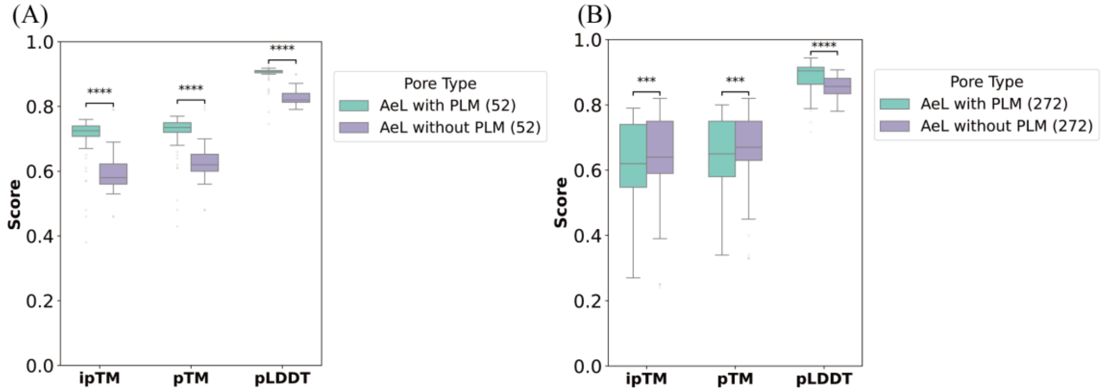


**Figure S3.** Comparison of multimer-level self-confidence scores (ipTM, pTM, and average pLDDT) for AeL models predicted by AF3 with and without PLM, respectively. (A) The subset of 52 models that switched conformations with addition of PLM molecules. (B) All 272 AF3 models. Significances of paired comparisons were calculated using two-sided Wilcoxon–Mann–Whitney U-test (****P* < 0.001, *****P* < 0.0001).

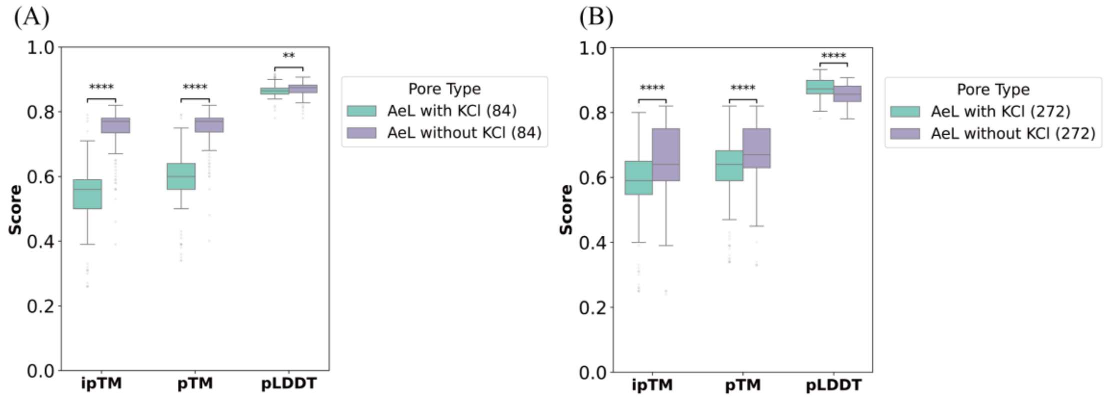


**Figure S4.** Comparison of multimer-level self-confidence scores (ipTM, pTM, and average pLDDT) for AeL models predicted by AF3 with and without $K^+$ and $Cl^-$ ions, respectively. (A) The subset of 84 models that switched conformations with addition of $K^+$ and $Cl^-$ ions. (B) All 272 AF3 models. Significances of paired comparisons were calculated using two-sided Wilcoxon–Mann–Whitney U-test (**$P < 0.01$, ****$P < 0.0001$).

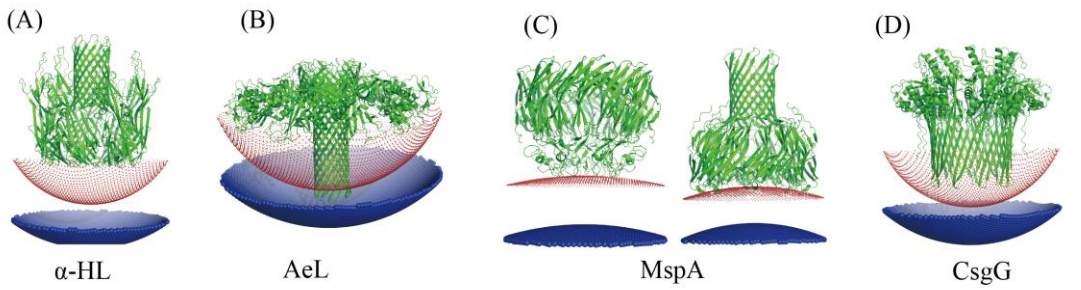


**Figure S5.** Examples of nanopore models unable to embed correctly. (A) α-HL (UniRef90 ID: UniRef90_UPI000D7CFDF9), (B) AeL (AFDB ID: A0A0T6TY63), (C) MspA (UniRef90 ID: UniRef90_A0A1D8T2P1, UniRef90 ID: UniRef90_UPI0027DB7C8D), and (D) CsgG (AFDB ID: A0A1T4SF25, AFDB ID: A0A4U2EGC6).

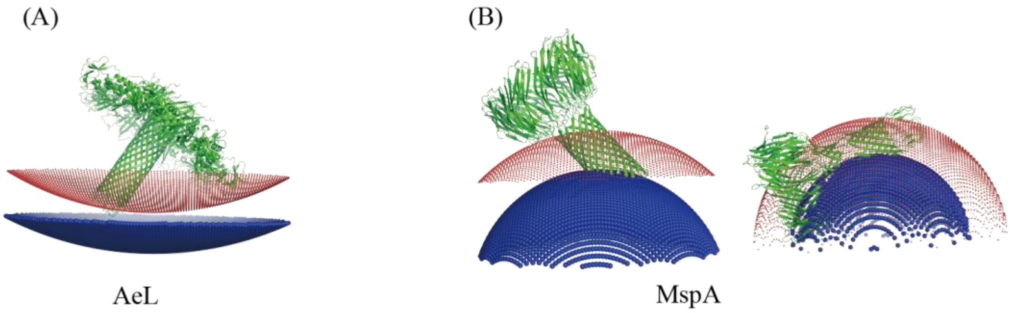


**Figure S6.** Examples of nanopore protein models with tilt angle > 10°. (A) AeL (UniRef90 ID: UniRef90_UPI00202CC0C0), (B) MspA (UniRef90 ID: UniRef90_UPI001CCE9F94, UniRef90 ID: UniRef90_UPI00326714A9).

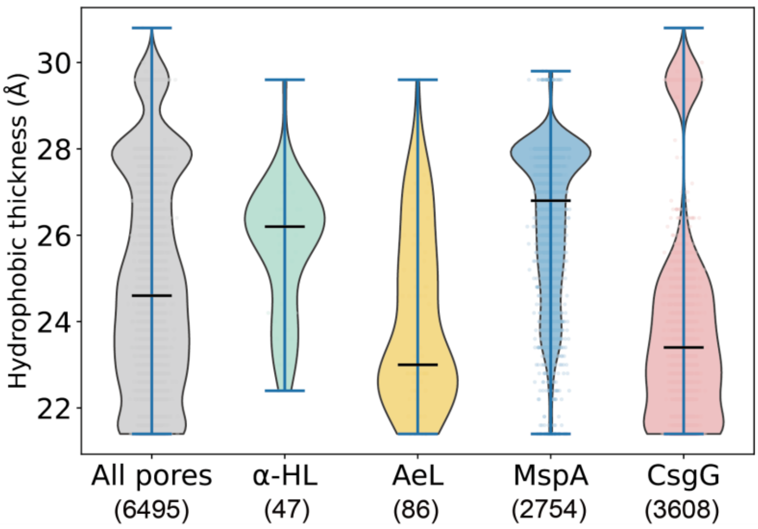

**Figure S7.** Violin plot of hydrophobic thickness (Å) for all nanopore candidates.

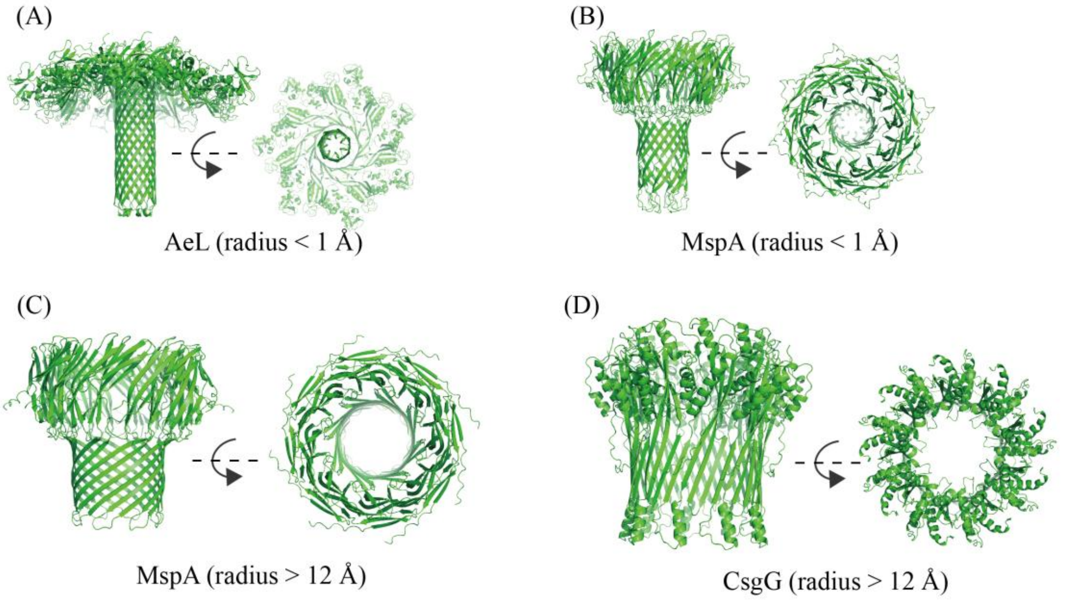


**Figure S8.** Examples of nanopore protein models with radius < 1 Å or > 12 Å. (A) AeL (radius = 0.98 Å, UniRef90 ID: UniRef90_A0A0F5VB76), (B) MspA (radius = 0.69 Å, AFDB ID: AF-A0A1A1WW01), (C) MspA (radius = 12.43 Å, UniRef90 ID: UniRef90_A0A3M2L262), and (D) CsgG (radius = 13.65 Å, AFDB ID: AF-A0A6M4GPJ4).

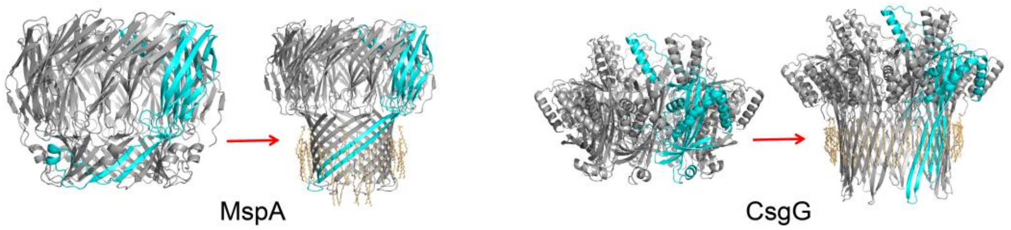


**Figure S9.** Examples of switching conformation by adding PLM in co-folding prediction of MspA (UniRef90 ID: UniRef90_A0A516WL96) and CsgG (AFDB ID: AF-A0A6N9RID5). The monomer chain of each protein is highlighted in cyan. PLM molecules are in orange.

**Table S1.** Experimentally resolved pore-like structures of representative nanopore types in the PDB.

| Nanopore | PDB ID (total number of representative experimental structures) |
|---|---|
| α-HL | 7AHL, 6U49, 3M2L, 3M3R, 7O1Q, 3M4E, 8JX2, 6U4P, 8JX3, 4P24, 3ANZ, 3M4D (12) |
| AeL | Prepore: 5JZH, 9FMX (2)<br>Post-prepore/quasipore: 5JZW (1)<br>Final pore: 5JZT, 6RB9, 9FNP, 9FNQ, 9FML, 9FM6 (6) |
| MspA | 1UUN (1) |
| CsgG | 6LQJ, 6L7C, 6LQH, 7BRM, 6L7A, 6SI7, 4UV3, 4Q79, 3X2R (9) |

**Table S2.** Model counts of each step of the mining workflow (step 1 to 5) and improved embedding in joint AF3-predictions with PLM molecules for incorrectly embedded AeL, MspA, and CsgG models.

| | α-HL | AeL | MspA | CsgG |
|---|---|---|---|---|
| Step1: Name search | 998 | 548 | 1712 | 6036 |
| Step2: Structure-based search | 39 | 293 | 1364 | 1179 |
| Step3: Sequence-based search | 75 | 61 | 1726 | 2466 |
| Step4: Multimeric structure prediction | 103 | 353 | 3026 | 3618 |
| Step5: High-quality structure filtration (AFM : AF3) | 47 (43 : 4) | 272 (36 : 236) | 2754 (2293 : 461) | 3608 (1072 : 2536) |
| Models embedded correctly (AFM : AF3) | 46 (42 : 4) | 20 (0 : 20) [1] | 2663 (2229 : 434) | 3118 (1030 : 2088) |
| Models embedded incorrectly (AFM : AF3) | 1 (1 : 0) | 66 (2 : 64) | 91 (64 : 27) | 490 (42 : 448) |
| Models embedded incorrectly with ill-predicted TM region [2] (AFM : AF3) | N.A. | N.A. | 32 (21 : 11) | 41 (9 : 32) |
| Models able to embed correctly with PLM (AFM : AF3) | N.A. | 31 (0 : 31) [3] | 26 (18 : 8) [4] | 19 (1 : 18) [4] |

Note:

1. Only AeL models in the final pore state were subjected to membrane embedding analysis, including 2 AFM models and 84 AF3 models (Figure 2D right).

2. Models embedded incorrectly and exhibiting visible structural distortions in the transmembrane (TM) region.

3. 31 out of 66 incorrectly embedded AeL models were able to embed correctly in the joint AF3 prediction with PLM molecules.

4. 26 out of 32 incorrectly embedded MspA models with structural distortion in TM region and 19 out of 41 CsgG models were able to embed correctly in the joint AF3 prediction with PLM molecules.

**Table S3.** Structural similarity (TM-score) of the experimental structures of AeL nanopores to reference conformations.

| PDB ID (Conformation) | Prepore | Post-prepore | Quasipore | Final pore |
|---|---|---|---|---|
| 5JZH (Prepore) | 1 | 0.950 | 0.691 | 0.666 |
| 9FMX (Prepore) | 1 | 0.951 | 0.691 | 0.667 |
| 5JZW (Post-prepore) | 0.858 | 1 | 0.709 | 0.674 |
| 5JZW (Quasipore) | 0.655 | 0.745 | 1 | 0.947 |
| 5JZT (Final pore) | 0.620 | 0.702 | 0.955 | 1 |
| 6RB9 (Final pore) | 0.266 | 0.322 | 0.461 | 1 |
| 9FM6 (Final pore) | 0.635 | 0.713 | 0.959 | 1 |
| 9FML (Final pore) | 0.664 | 0.744 | 0.985 | 1 |
| 9FNP (Final pore) | 0.615 | 0.698 | 0.942 | 1 |
| 9FNQ (Final pore) | 0.627 | 0.710 | 0.957 | 1 |

Note: The prepore state of AeL has two experimental structures and the final pore states has six. The structure similarity of a structure to a given conformational state refers to the highest TM-score of that structure compared to all the structures belonging to the state.

**Table S4.** AF3 prediction results of AeL-like nanopores with ligands.

| | AeL | AeL + PLM [1] | AeL + 10 $K^+$ + 10 $Cl^-$ |
|---|---|---|---|
| Prepore | 183 | 140 | 231 |
| Post-prepore | 1 | 0 | 0 |
| Quasipore | 0 | 0 | 0 |
| Final pore | 84 | 127 | 22 |
| Other | 4 | 5 | 19 |
| Conformational change[2] | | 52 in total<br>**prepore to final pore: 46**<br>prepore to other: 1<br>post-prepore to final pore: 1<br>final pore to prepore: 4 | 84 in total<br>prepore to final pore: 4<br>prepore to other: 13<br>post-prepore to prepore: 1<br>**final pore to prepore: 64**<br>final pore to other: 2 |
| Final pore able to embed | 20 | 68 in total<br>45 kept to be final pore<br>23 converted from prepore | N.A. |
| Final pore able to embed correctly | 20 | 65 in total<br>42 kept to be final pore<br>23 converted from prepore | N.A. |

Note:

1. Number of added PLM molecules were determined based the number of available tokens by AF3 online server and ranged from 36 to 54 (40 for 272 AeL predictions on average) (methods).

2. Number of conformational transitions were referenced to the results of predicting the protein nanopore alone.
